# A chromosome-level genome assembly of “a living fossil”, the tadpole shrimp *Lepidurus arcticus* (Pallas, 1793)

**DOI:** 10.64898/2026.06.04.730062

**Authors:** Marius A Strand, Ole K Tørresen, Jens Ådne R Haga, Bram Danneels, Morten Skage, Giada Ferrari, Ave Tooming-Klunderud, Dag O Hessen, Kjetill S Jakobsen

## Abstract

We present the first chromosome-level reference genome for *Lepidurus arcticus* (Pallas, 1793), a freshwater crustacean with circumpolar distribution. *L. arcticus* belongs to the small order of freshwater Notostracan crustaceans that are representatives of the ancient group Branchiopoda. This group has a remarkable morphological stability and is frequently labelled “living fossils”. Its ancient origin, streamlined genome (estimated to 0.11 Gb) and reproductive flexibility makes this a very interesting candidate for genomic studies.

The haplotype-resolved assemblies are composed of two pseudo-haplotypes spanning 81.2 megabases (Mb) and 81.8 Mb, respectively, and each scaffolded into 6 chromosomes. Both haplotypes (hap) show high completeness and identical BUSCO scores of 98.3 for hap1 and hap2. The scaffold N50 length is 13.4 Mb for hap1 and 13.9 Mb for hap2, and k-mer completeness estimated from PacBio HiFi reads was 95.79% and 96.18%, respectively. The haplotypes display very low estimated genome-wide heterozygosity of 0.133%. The assembly contains 10901 (hap1) and 10910 (hap2) protein-coding genes. Repetitive elements comprised approximately 24–25% of each haplotype, with long terminal repeat retrotransposons representing the most abundant transposable element class at approximately 8–9%.

Comparison with the near chromosome-level genome of *Lepidurus packardi* revealed substantial intrachromosomal rearrangements, despite similar chromosome numbers and chromosome sizes. Differences in transposable element content between *L. arcticus* and *L. packardi* were primarily driven by retrotransposons, particularly LTR and LINE elements. This reference genome provides a valuable resource for future population genomic studies and for investigating evolutionary stasis at the genome level.

## Introduction

*Lepidurus arcticus* (Pallas, 1793) belongs to the small order of freshwater notostracan crustaceans that are representatives of the ancient group Branchiopoda. Judged from morphological criteria, this is an evolutionary remarkably stable order that has existed since the Permian > 250 million years ago, and the earliest notostracan dates back to the upper Devonian (approximately 420–360 million years ago; Lagebro et al. (2015). Fossil records dating more than 200 million years back to the Triassic are known with striking similarity of present members of these genera (Longhurst, 1955; Mathers, Hammond, Jenner, Hänfling, et al., 2013; Tasch, 1969). These lineages are sometimes referred to as examples of evolutionary stasis or ‘living fossils’ (King & Hanner, 1998; Longhurst, 1958; Mantovani et al., 2004).

Notostracans inhabit a highly diverse range of habitats including harsh ecological systems such as desert pools and alpine or high Arctic ponds, with *L. arcticus* as a “cold stenotherm” species. This species has a distinctive northern, circumpolar distribution and may occur both in shallow moraine ponds and large lakes in arctic, subarctic or alpine areas (Hessen et al., 2004; Qvenild et al., 2025). During the Pleistocene glaciations, the species was apparently far more widespread in western Europe than presently (Bennie, 1894; Longhurst, 1955).

Despite general morphological stability over geological periods, notostracans possess a phenotypic flexibility within the major morphological frame that renders taxonomic affinities difficult (Hessen et al., 2004; King & Hanner, 1998; Mathers, Hammond, Jenner, Zierold, et al., 2013). The apparent long-term evolutionary stasis could be explained by stabilizing selection, but the strong morphological flexibility may indicate a high level of heterozygosity. Not the least is the reproductive flexibility quite remarkable among the Notostracans, with transitions between sexual and asexual reproduction, as well as the very rare reproductive mode of androdioecy – a sexual system where males and hermaphrodites co-occur in varying frequencies in populations, with different levels of self-fertilisation and outcrossing (Mathers, Hammond, Jenner, Zierold, et al., 2013; Weeks et al., 2006).

Studies of North American species of *Lepidurus* based on allozyme studies and sequencing of a 330 bp 12S rDNA subunit confirmed the presence of five distinct species, yet with a rather pronounced macrogeographical overlap (King & Hanner, 1998). This study did, however, not include the most northern species, *L. arcticus*. A circumpolar assessment of *L. arcticus* based on mitochondrial sequencing from 48 populations revealed two distinct haplogroups, each consisting of separate haplotypes (Hessen et al., 2004). Alignment with published sequences of other species of *Lepidurus* clearly revealed that all the studied populations were distinctly different from other species.

The distribution of *L. arcticus* has changed in response to climate and ice coverage of northern latitude, but in the recent millennia it has likely experienced a strong reduction due to extensive fish stocking, and in the recent decades elevated temperatures pose a risk to this species (Qvenild et al., 2025) as well as other ancient and rare taxa of branchiopods (Lindholm et al., 2015). Both its ancient origin, its remarkable morphological conservation alongside a genomic and reproductive flexibility as well as its existence in extreme and marginal habitats undergoing rapid changes to the extensive warming at high latitudes, makes this species a highly interesting candidate for sequencing and genomic analysis. Like all branchiopods it has a small genome (Alfsnes et al., 2017), around 0.11 pg (∼0.108 Gb) (Savojardo et al., 2019), making it a good candidate for sequencing and assembly, yet there are few such attempts. Savojardo et al. (2019) sequenced *L. arcticus* and *Lepidurus apus lubbocki* where 73.2 Mb (*L. arcticus*) and 90.3 Mb (*L. apus lubbocki*) were assembled, respectively. The assemblies covered up to 84% of the estimated genome size, and revealed that 13%–16% of the assembled genomes consisted of repeats and that there were some differences in their transposable element content.

A near chromosome-level reference genome exists only for *Lepidurus packardi* (Kieran Blair et al., 2022), an endemic and rare species inhabiting vernal ponds in California, susceptible to habitat destruction and drought. They assembled a genome of 108.6 Mb, with 6 chromosome-length scaffolds that comprised 71% of total genomic length and 444 total contigs. In total, they predicted 17,650 genes for this species. The estimated divergence time between *L. packardi, L. apus lubbocki* and *L. arcticus* is some 65 million years (Mathers, Hammond, Jenner, Hänfling, et al., 2013), and the species inhabit totally different habitats with no geographic overlap.

We here present the first chromosome-level reference genome for *L. arcticus* from the mountain plateau, Hardangervidda in Norway using long read sequencing (PacBio HiFi) and Hi-C for chromosome scaffolding. To date, this is the most complete *Lepidurus* reference genome that has been generated.

## Material and Methods

### Sample acquisition and DNA extraction

The sampling site (60.403196° N, 7.369452° E; 832 m a.s.l.) is a shallow, fishless, subalpine pond (∼1000 m^2^, <2 m depth) (Fig. 1) located in Eidfjord, Vestland, Norway in the outskirts of the Hardangervidda mountain plateau. Sampling was conducted by hand on 21 August 2024. Three adult individuals were transported alive to Oslo for sequencing in a 2 L temperature-controlled container filled with pond water. One specimen was selected for genome sequencing: head tissue was used for PacBio HiFi sequencing, and thorax and abdomen from the same specimen were used for Hi-C library preparation.

**Figure 1.**
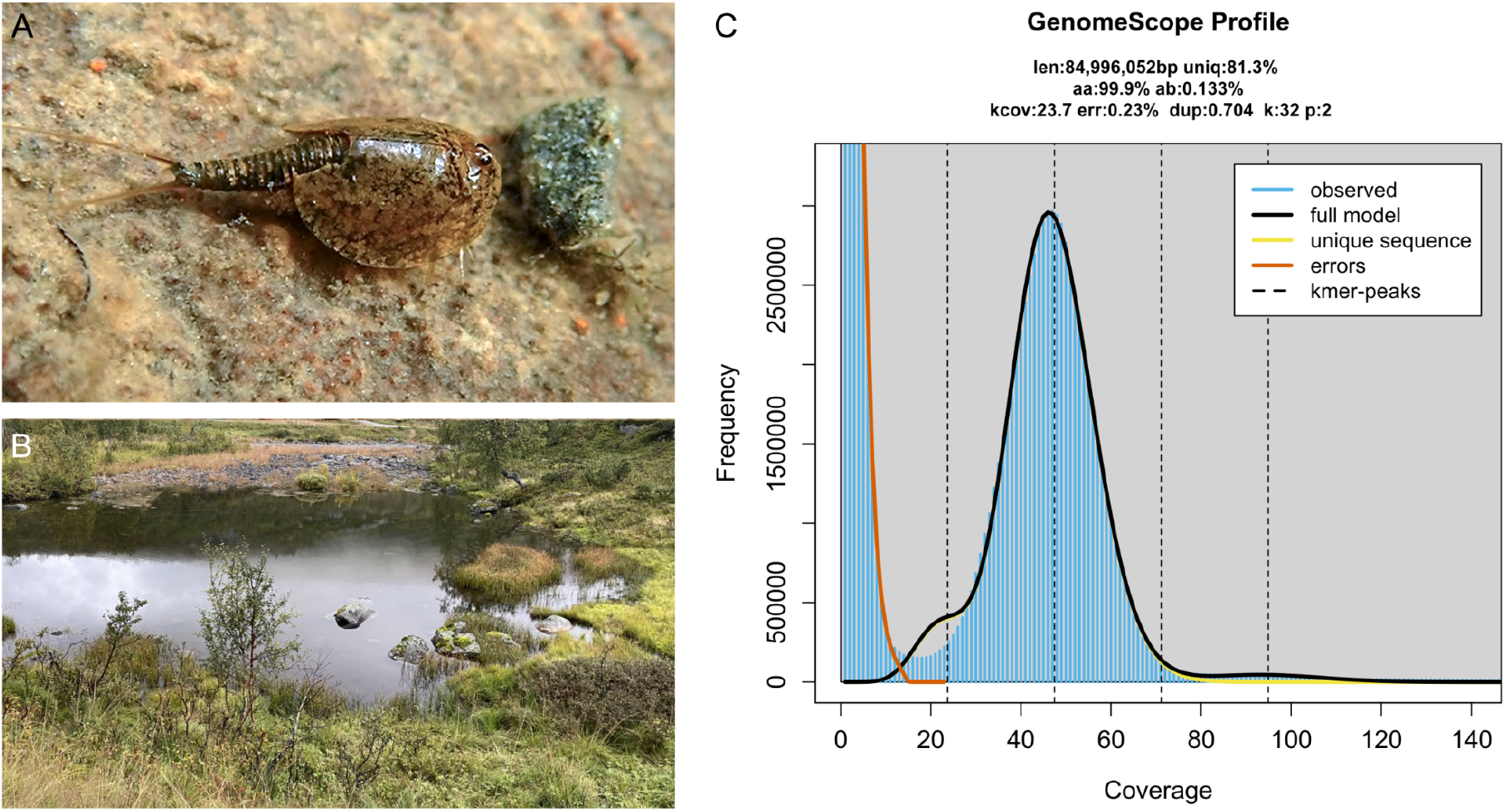
Sequenced specimen and genome profile. **A)** Representative photograph of *L. arcticus* from the same pond as the sequenced individual; photo by Jens Ådne R. Haga. **B)** Collection environment of the sequenced *L. arcticus* specimen; photo by Jens Ådne R. Haga. **C)** GenomeScope profile of the HiFi reads from the sequenced individual. This analysis estimates a 85 Mb genome, with 0.133% heterozygosity. The left-hand peak of k-mers (appearing as a shoulder) corresponds to k-mers from heterozygous regions of the genome, while the right-hand peak is from homozygous regions.

HMW DNA for PacBio long read sequencing was isolated from head tissue using Genomic Tip (Qiagen) according to manufacturer’s recommendations. Quality check of amount, purity and integrity of isolated DNA was performed using Qubit BR DNA quantification assay kit (Thermo Fisher), Nanodrop (Thermo Fisher), and Fragment Analyser (DNA HS 50kb large fragment kit, Agilent Tech.).

### Library preparation and sequencing for *de novo* assembly

Before PacBio HiFi library preparation, DNA was purified an additional time using AMPure PB beads (1:1 ratio). Purified HMW DNA was sheared into an average fragment size of 15-20 kbp large fragments using the Megaruptor3 (Diagenode). A HiFi library was prepared following the PacBio protocol for HiFi library preparation using the SMRTbell® Prep Kit 3.0. The final HiFi library was size-selected with a 9 kbp cut-off using a BluePippin (Sage Sciences) and sequencing was performed by the Norwegian Sequencing Centre on a PacBio Revio instrument using the SPRQ chemistry (Pacific Biosciences Inc), generating 11.7 Gb of HiFi data.

A Hi-C library was prepared using the Arima High Coverage HiC kit (Arima Genomics), following the low input protocol in the User Guide for Animal Tissues (document part number A160162 v01) using thorax and abdomen of the same specimen. Final library quality was quantified using a Kapa Library quantification kit for Illumina (Roche Inc.). The library was sequenced with other libraries on a NovaSeq X instrument (Illumina Inc) with 2 × 150 bp paired end mode at the Norwegian Sequencing Centre (https://www.sequencing.uio.no).

### Genome assembly and curation, annotation and evaluation

A full list of relevant software tools and versions is presented in Table 1. We assembled the species using a pre-release of the EBP-Nor genome assembly pipeline (https://github.com/ebp-nor/GenomeAssembly). KMC (Kokot et al., 2017) was used to count k-mers of size 32 in the PacBio HiFi reads, excluding k-mers occurring more than 10,000 times. GenomeScope (Ranallo-Benavidez et al., 2020) was run as part of the pipeline on the k-mer histogram output from KMC and was included in the methods for completeness. Ploidy level was investigated using Smudgeplot (Ranallo-Benavidez et al., 2020). HiFiAdapterFilt (Sim et al., 2022) was applied on the HiFi reads to remove possible remnant PacBio adapter sequences. The filtered HiFi reads were assembled using Hifiasm (Cheng et al., 2021) with Hi-C integration resulting in a pair of haplotype-resolved assemblies, pseudo-haplotype one (qbLepArct1.1.hap1, hereby hap1) and pseudo-haplotype two (qbLepArct1.1.hap2, hereby hap2). Unique k-mers in each assembly/pseudo-haplotype were identified using meryl (Rhie et al., 2020) and used to create two sets of Hi-C reads, one without any k-mers occurring uniquely in hap1 and the other without k-mers occurring uniquely in hap2.

**Table 1.**
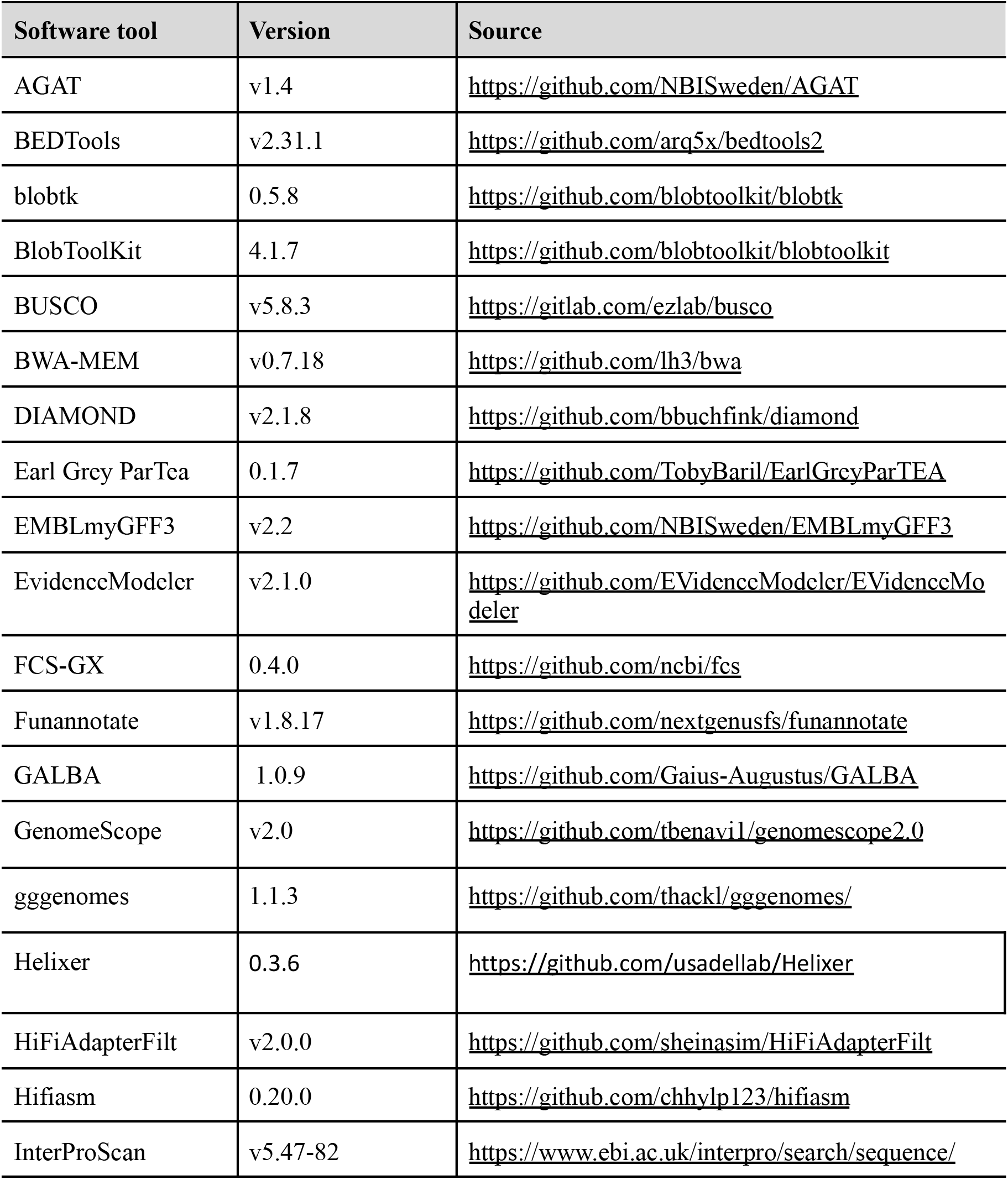
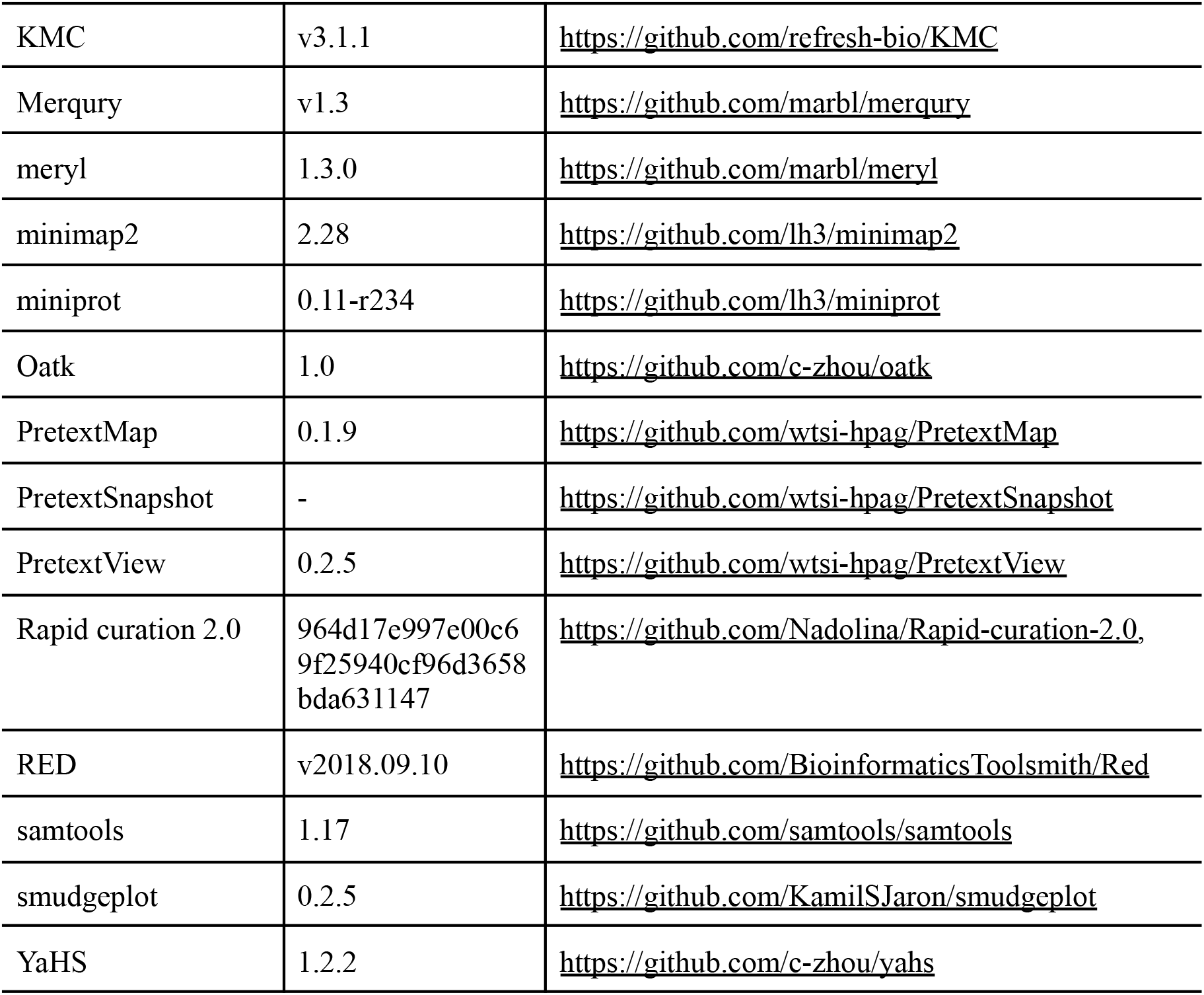
Software tools: versions and sources.

K-mer filtered Hi-C reads were aligned to each scaffolded assembly using BWA-MEM (Li, 2013) with -5SPM options. The alignments were sorted based on name using samtools (Li et al., 2009) before applying samtools fixmate to remove unmapped reads and secondary alignments and to add mate score, and samtools markdup to remove duplicates. The resulting BAM files were used to scaffold the two assemblies using YaHS (Zhou et al., 2023) with default options. FCS-GX (Astashyn et al., 2024), BlobToolKit and BlobTools2 (Laetsch & Blaxter, 2017) were used to search for putative contamination and contaminated sequences were removed. BlobToolKit coverage-GC plots were generated to inspect scaffold-level coverage, GC content, and taxonomic assignment (Supplementary Fig. 2). The mitochondrion was searched for in contigs and reads using Oatk (Zhou et al., 2024). The assemblies were manually curated using PretextView and Rapid curation 2.0. Chromosomes were identified by inspecting the Hi-C contact map in PretextView.

We annotated the genome assemblies using a pre-release version of the EBP-Nor genome annotation pipeline (https://github.com/ebp-nor/GenomeAnnotation). First, AGAT (https://zenodo.org/record/7255559) agat_sp_keep_longest_isoform.pl and agat_sp_extract_sequences.pl were used on the *Daphnia pulex* (GCF_021134715.1) genome assembly and annotation to generate one protein (the longest isoform) per gene. Miniprot (Li, 2023) was used to align the proteins to the curated assemblies. UniProtKB/Swiss-Prot (UniProt Consortium, 2023) release 2025_03 in addition to the Arthropoda part of OrthoDB v12 (Tegenfeldt et al., 2025) were also aligned separately to the assemblies. RED (Girgis, 2015) was run via redmask (https://github.com/nextgenusfs/redmask) on the assemblies to mask repetitive areas. In addition to gene models from protein alignments, GALBA (Brůna et al., 2023; Buchfink et al., 2015; Hoff & Stanke, 2019; Li, 2023; Stanke et al., 2006) was run with the *Daphnia pulex* proteins using the miniprot mode on the masked assemblies to generate *ab initio* predicted gene models. Helixer(Holst et al., 2025) was run using the invertebrate-specific model (invertebrate_v0.3_m_0100).

The funannotate-runEVM.py script from Funannotate was used to run EvidenceModeler (Haas et al., 2008) on the alignments of *Daphnia pulex* proteins to the *Lepidurus arcticus* genome assembly, UniProtKB/Swiss-Prot proteins, Arthropoda proteins from OrthoDB, and the predicted genes from GALBA and Helixer. The resulting predicted proteins were compared to the protein repeats that Funannotate distributes, using DIAMOND blastp, and the predicted genes were filtered based on this comparison using AGAT. The filtered proteins were compared to the UniProtKB/Swiss-Prot release 2025_03 using DIAMOND (Buchfink et al., 2015) blastp to find gene names and InterProScan (Jones et al., 2014) was used to discover functional domains. AGAT’s agat_sp_manage_functional_annotation.pl was used to attach the gene names and functional annotations to the predicted genes. EMBLmyGFF3 (Norling et al., 2018) was used to combine the FASTA files and GFF3 files into EMBL format for submission to ENA.

All the evaluation tools have also been implemented in a pipeline, similar to assembly and annotation (https://github.com/ebp-nor/GenomeEvaluation). Merqury (Rhie et al., 2020) was used to assess the completeness and quality of the genome assemblies by comparing them to the k-mer content of both the Hi-C reads and PacBio HiFi reads. BUSCO (Manni et al., 2021) was used to assess the completeness of the genome assemblies by comparing against the expected gene content in the arthropoda lineage. Gfastats (Formenti et al., 2022) was used to output different assembly statistics of the assemblies.

BlobToolKit and BlobTools2 (Laetsch & Blaxter, 2017), in addition to blobtk were used to visualize assembly statistics. To generate the Hi-C contact map image, the Hi-C reads were mapped to the assemblies using BWA-MEM (Li, 2013) using the same approach as above. Finally, PretextMap was used to create a contact map which was visualized using PretextSnapshot.

To assess whole-genome synteny between *L. arcticus* and *L. packardi*, we generated pairwise assembly alignments. Alignments were generated between the *L. packardi* assembly LEPA1 (GCA_023053545) (Kieran Blair et al., 2022) and our pseudo-haplotype assemblies with minimap2 using the (-x asm10) (Li, 2018). For *L. packardi*, we treated PGA_scaffold1-6 as putative homologous chromosomes based on their size and synteny relative to our curated *L. arcticus* assemblies. For visualization purposes, some large *L. packardi* scaffolds were reverse-complemented to match the orientation of the *L. arcticus* assemblies. Alignments with mapping quality <30 or alignment length <50 kb were excluded, and the retained alignments were visualized in R using gggenomes (Hackl et al., 2024).

Transposable elements were annotated for all assemblies using Earl Grey ParTEA, which extends TE annotation to multi-genome comparative analyses (Baril et al., 2024). Repeat landscapes were generated from the filtered repeat-annotation GFF files (*.filteredRepeats.gff). Fully nested Earl Grey annotations (NESTED=FULLY_NESTED) were removed, and repeat coverage was scaled by assembly size. Kimura two-parameter distances were extracted from the GFF attributes and binned to estimate the proportion of each assembly covered by repeat copies at different divergence levels. Repeat annotations were grouped into broad classes: LTR, LINE, SINE, DNA transposons, rolling-circle elements, other elements, and unclassified elements. The repeat landscape plot was generated with R.

To compare repeat distributions along chromosomes, analyses were restricted to the six largest scaffolds from each assembly. Scaffold lengths were calculated from the corresponding assembly FASTA files, and each scaffold was divided into non-overlapping 500 kb windows. Repeat coverage was calculated as the fraction of each window covered by each repeat class. Coverage values were normalized where overlapping annotations caused summed coverage to exceed the window length. Window-based repeat summaries were generated with R.

TE coverage was calculated from non-nested annotations after merging overlapping intervals. Overlap was calculated as the difference between summed annotation length and merged coverage. Fully nested annotations were collapsed and measured separately as unique genomic coverage. Reported TE coverage (Supplementary Table 1) therefore represents non-nested, non-overlapping repeat sequence, with overlap and nested fractions shown separately.

Part of the text in Methods and Results is based on a template we use for all the species we publish in the EBP-Nor project.

## Results

### *De novo* genome assembly and annotation

The genome of *Lepidurus arcticus* was assembled from a total of 144-fold coverage in Pacific Biosciences single-molecule HiFi long reads and 644-fold coverage in Arima Hi-C reads resulting in two haplotype-separated assemblies. The final assemblies have total lengths of 81.2 Mb and 81.8 Mb (Table 2 and Figure 2), respectively. Pseudo-haplotypes one (hap1) and two (hap2) have scaffold N50 size of 13.4 Mb and 13.9 Mb, respectively, and contig N50 of 12.8 Mb and 13.9 Mb, respectively (Table 2 and Fig. 2). Six pseudo-chromosomes were identified in both pseudo-haplotypes, as demonstrated by the Hi-C contact maps (Fig. 3).

**Table 2.**
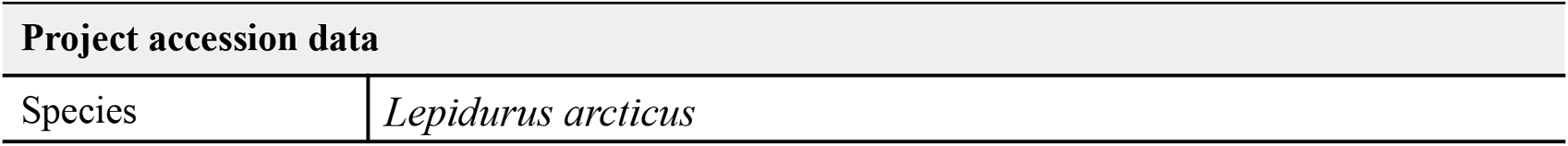
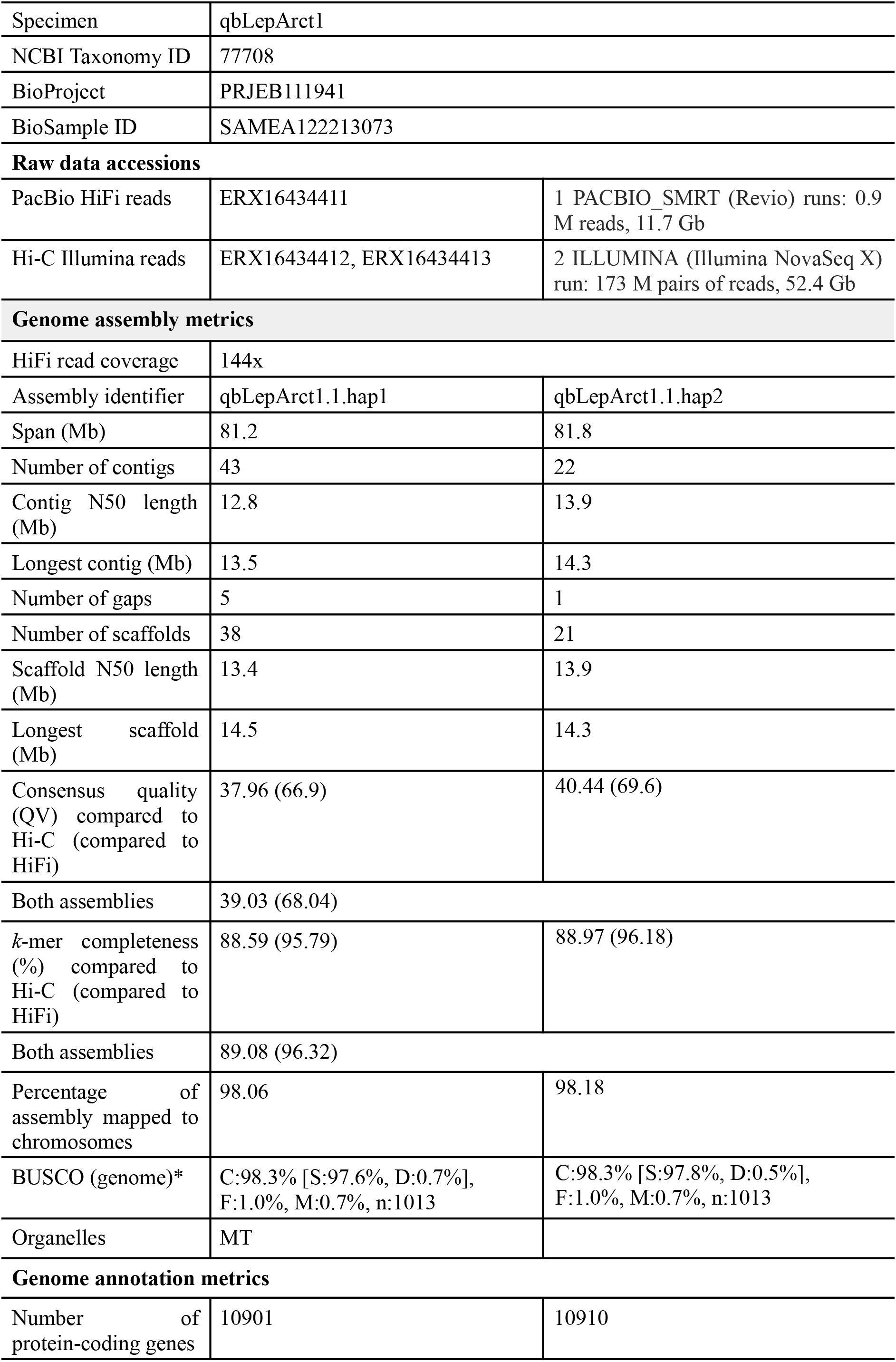
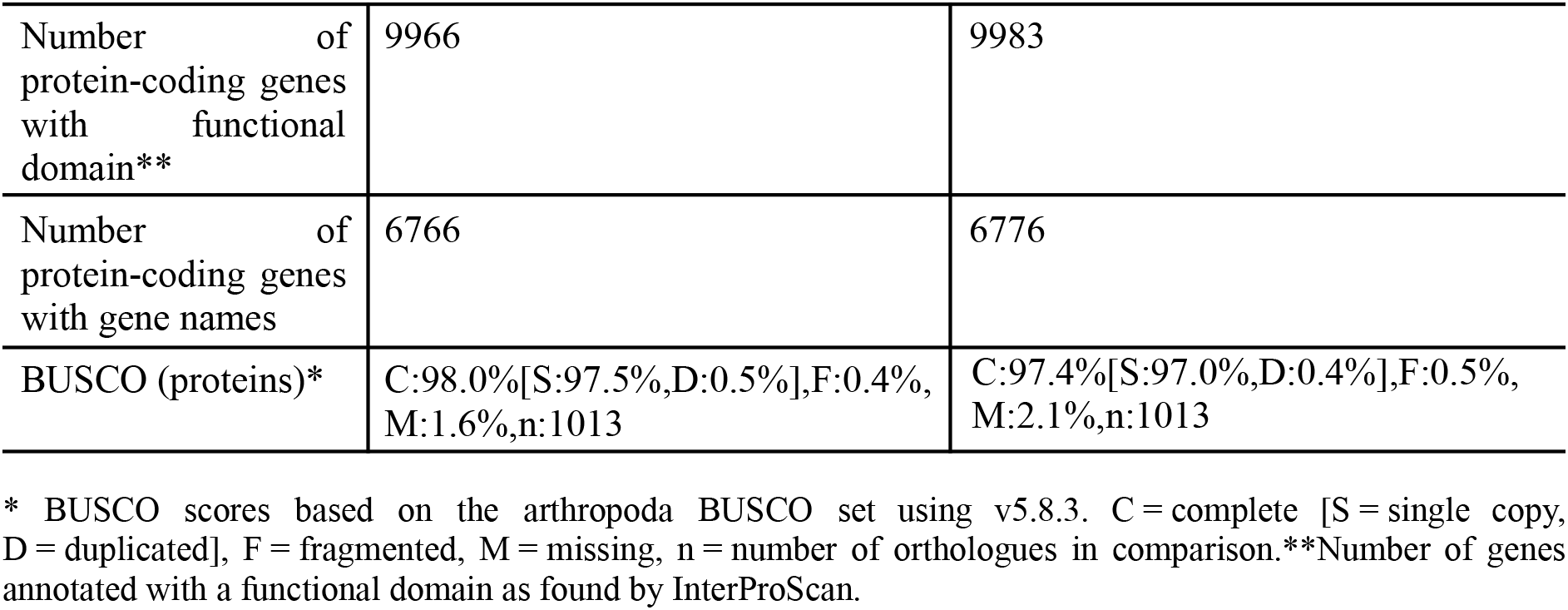
Genome data for *Lepidurus arcticus*.

| Project accession data |  |
| --- | --- |
| Species | *Lepidurus arcticus* |

|  |  |  |
| --- | --- | --- |
| Specimen | qbLepArct1 |  |
| NCBI Taxonomy ID | 77708 |  |
| BioProject | PRJEB111941 |  |
| BioSample ID | SAMEA122213073 |  |
| Raw data accessions |  |  |
| PacBio HiFi reads | ERX16434411 | 1 PACBIO_SMRT (Revio) runs: 0.9 M reads, 11.7 Gb |
| Hi-C Illumina reads | ERX16434412, ERX16434413 | 2 ILLUMINA (Illumina NovaSeq X) run: 173 M pairs of reads, 52.4 Gb |
| Genome assembly metrics |  |  |
| HiFi read coverage | 144x |  |
| Assembly identifier | qbLepArct1.1.hap1 | qbLepArct1.1.hap2 |
| Span (Mb) | 81.2 | 81.8 |
| Number of contigs | 43 | 22 |
| Contig N50 length (Mb) | 12.8 | 13.9 |
| Longest contig (Mb) | 13.5 | 14.3 |
| Number of gaps | 5 | 1 |
| Number of scaffolds | 38 | 21 |
| Scaffold N50 length (Mb) | 13.4 | 13.9 |
| Longest scaffold (Mb) | 14.5 | 14.3 |
| Consensus quality (QV) compared to Hi-C (compared to HiFi) | 37.96 (66.9) | 40.44 (69.6) |
| Both assemblies | 39.03 (68.04) |  |
| k-mer completeness (%) compared to Hi-C (compared to HiFi) | 88.59 (95.79) | 88.97 (96.18) |
| Both assemblies | 89.08 (96.32) |  |
| Percentage of assembly mapped to chromosomes | 98.06 | 98.18 |
| BUSCO (genome)* | C:98.3% [S:97.6%, D:0.7%], F:1.0%, M:0.7%, n:1013 | C:98.3% [S:97.8%, D:0.5%], F:1.0%, M:0.7%, n:1013 |
| Organelles | MT |  |
| Genome annotation metrics |  |  |
| Number of protein-coding genes | 10901 | 10910 |
| Number of protein-coding genes with functional domain** | 9966 | 9983 |
| Number of protein-coding genes with gene names | 6766 | 6776 |
| BUSCO (proteins)* | C:98.0%[S:97.5%,D:0.5%],F:0.4%,M:1.6%,n:1013 | C:97.4%[S:97.0%,D:0.4%],F:0.5%,M:2.1%,n:1013 |

\* BUSCO scores based on the arthropoda BUSCO set using v5.8.3. C = complete [S = single copy, D = duplicated], F = fragmented, M = missing, n = number of orthologues in comparison. \*\*Number of genes annotated with a functional domain as found by InterProScan.

**Figure 2.**
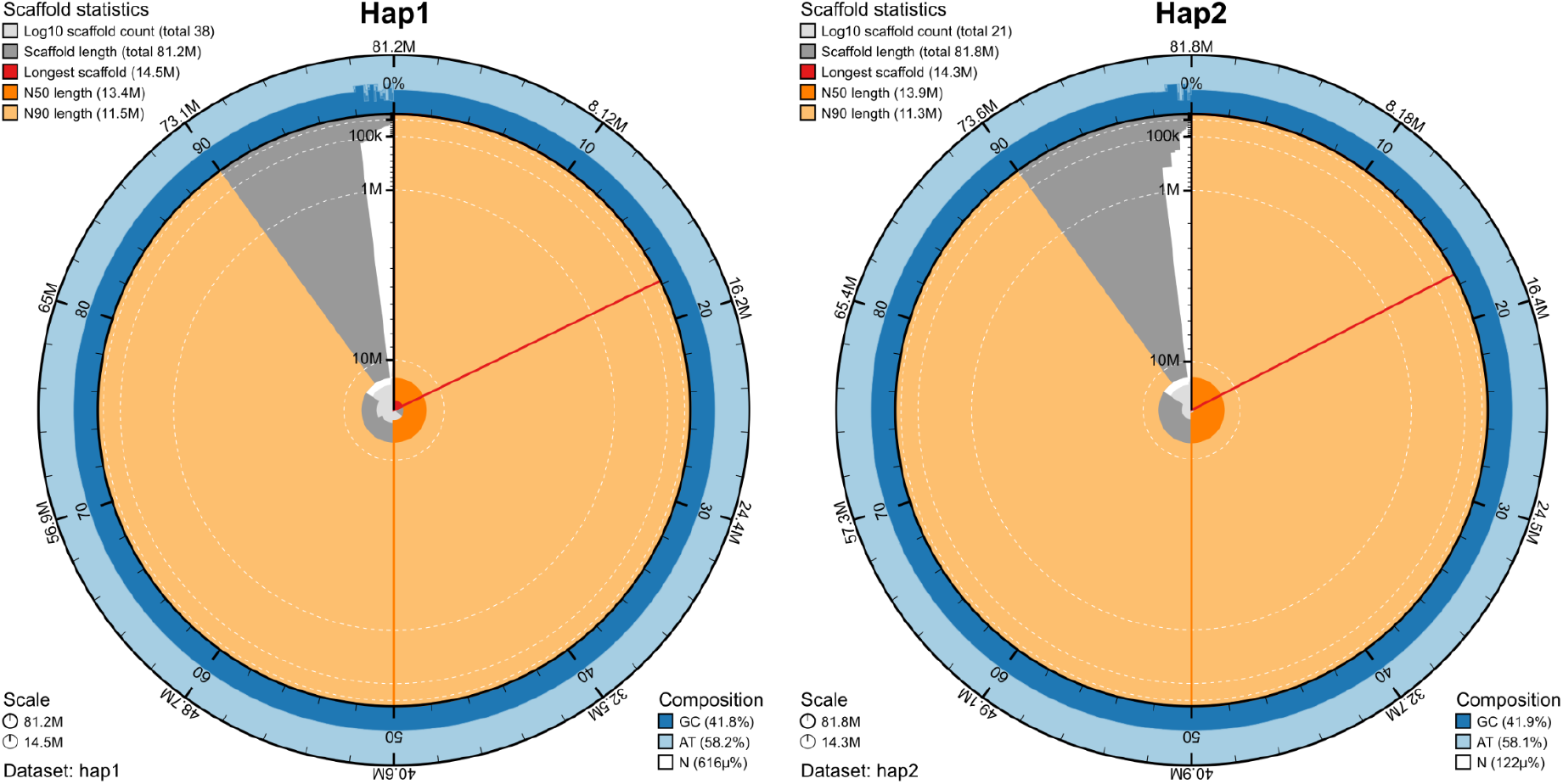
Metrics of the genome assemblies of *Lepidurus arcticus* hap1 and hap2. The BlobToolKit Snailplots show N50 metrics. The two outermost bands of the circle signify GC versus AT composition at 0.1% intervals, with mean, maximum, and minimum. The third outermost shows the N90 scaffold length, while the fourth is N50 scaffold length. The line from middle to second outermost band shows the size of the largest scaffold. All the scaffolds are arranged in a clockwise manner from largest to smallest, and shown in darker gray with white lines at different orders of magnitude, shown as a scale on the radius. The light gray shows the cumulative scaffold count. The scale inset in the lower left corner shows the total amount of sequence in the whole circle, and the fraction of the circle encompassed in the largest scaffold.

**Figure 3.**
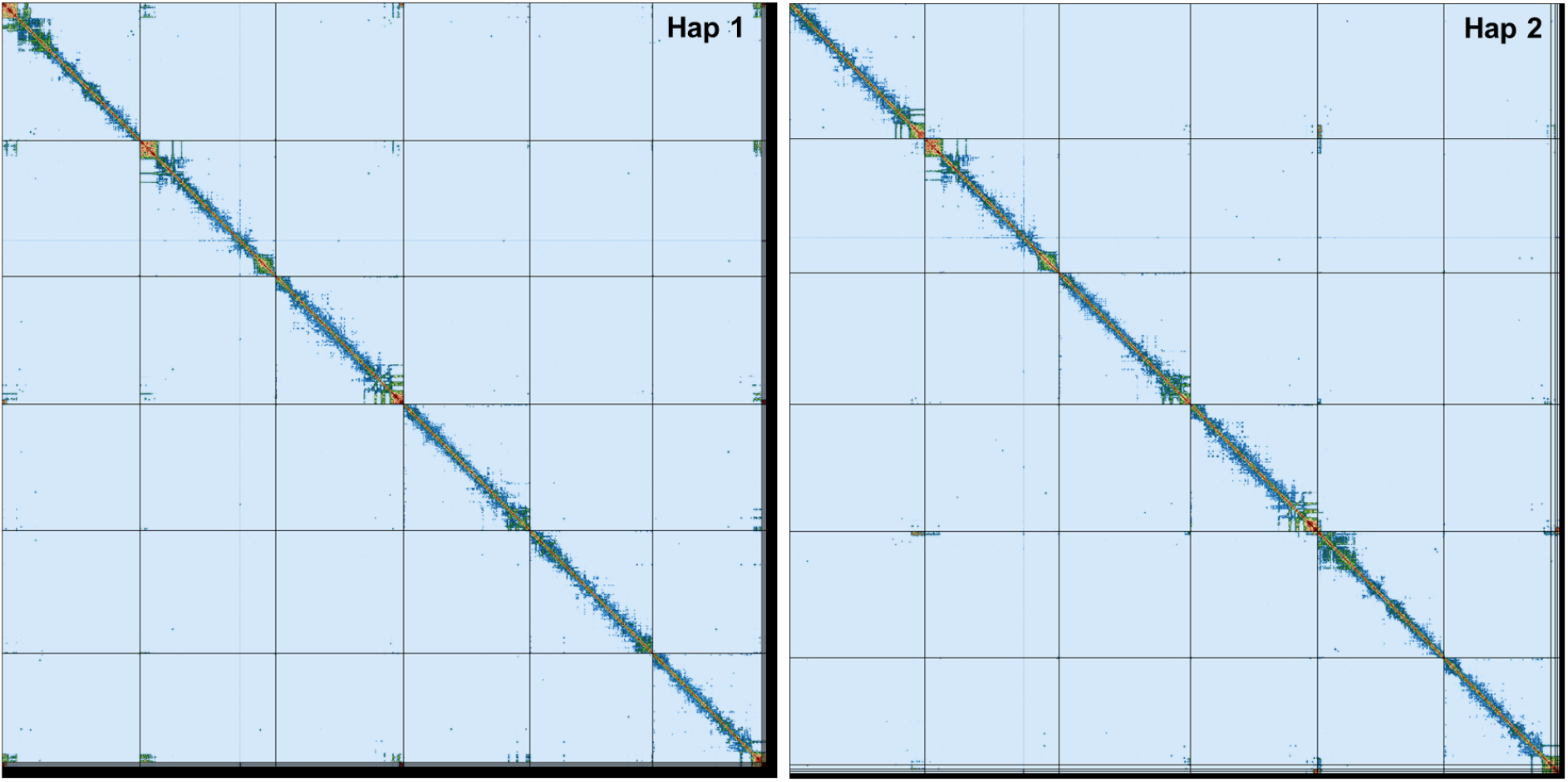
Hi-C contact maps for *Lepidurus arcticus* hap1 and hap2. The contact map displays interaction frequencies between genomic regions, where darker shades represent a higher number of Hi-C contacts. The axes correspond to the coordinates along each assembly. The Hi-C contact maps were generated by mapping the Hi-C reads to the haplotype genomes using BWA-MEM, generating a contact map using PretextMap and visualized using PretextSnapshot.

When compared to a k-mer database of the Hi-C reads, hap1 had a k-mer completeness of 88.6%, hap2 of 89.0%, and combined they have a completeness of 89.1% (Table 2). Further, hap1 had an assembly consensus quality value (QV) of 38.0 and hap2 of 40.44, where a QV of 40 corresponds to one error every 10,000 bp, or 99.99% accuracy compared to a k-mer database of the Hi-C reads (QV 66.9 and 69.6, respectively, compared to a k-mer database of the HiFi reads) (Table 2). Merqury k-mer spectra showed that most assembly k-mers were shared between the two pseudo-haplotypes, with only small haplotype-specific fractions, consistent with the low heterozygosity and high similarity between haplotypes (Supplementary Fig. 1). Coverage-GC profiles showed no major scaffold outliers with divergent GC content or coverage (Supplementary Fig. 2). A total of 10901 and 10910 protein-coding genes were annotated in hap1 and hap2, respectively (Table 2).

Whole-genome synteny analysis showed strong macrosynteny between *Lepidurus packardi* LEPA1 and both *L. arcticus* pseudo-haplotype assemblies, with the six largest LEPA1 scaffolds corresponding closely to the six curated *L. arcticus* chromosomes (Fig. 4). The two *L. arcticus* pseudo-haplotypes were broadly collinear, differing only by three small translocated segments and showing no major large-scale structural differences. Interchromosomal rearrangement between LEPA1 and *L. arcticus* appeared limited. In contrast, all six chromosomes showed intrachromosomal rearrangements, including inversions and local translocations that disrupt collinearity within chromosomes.

**Figure 4.**
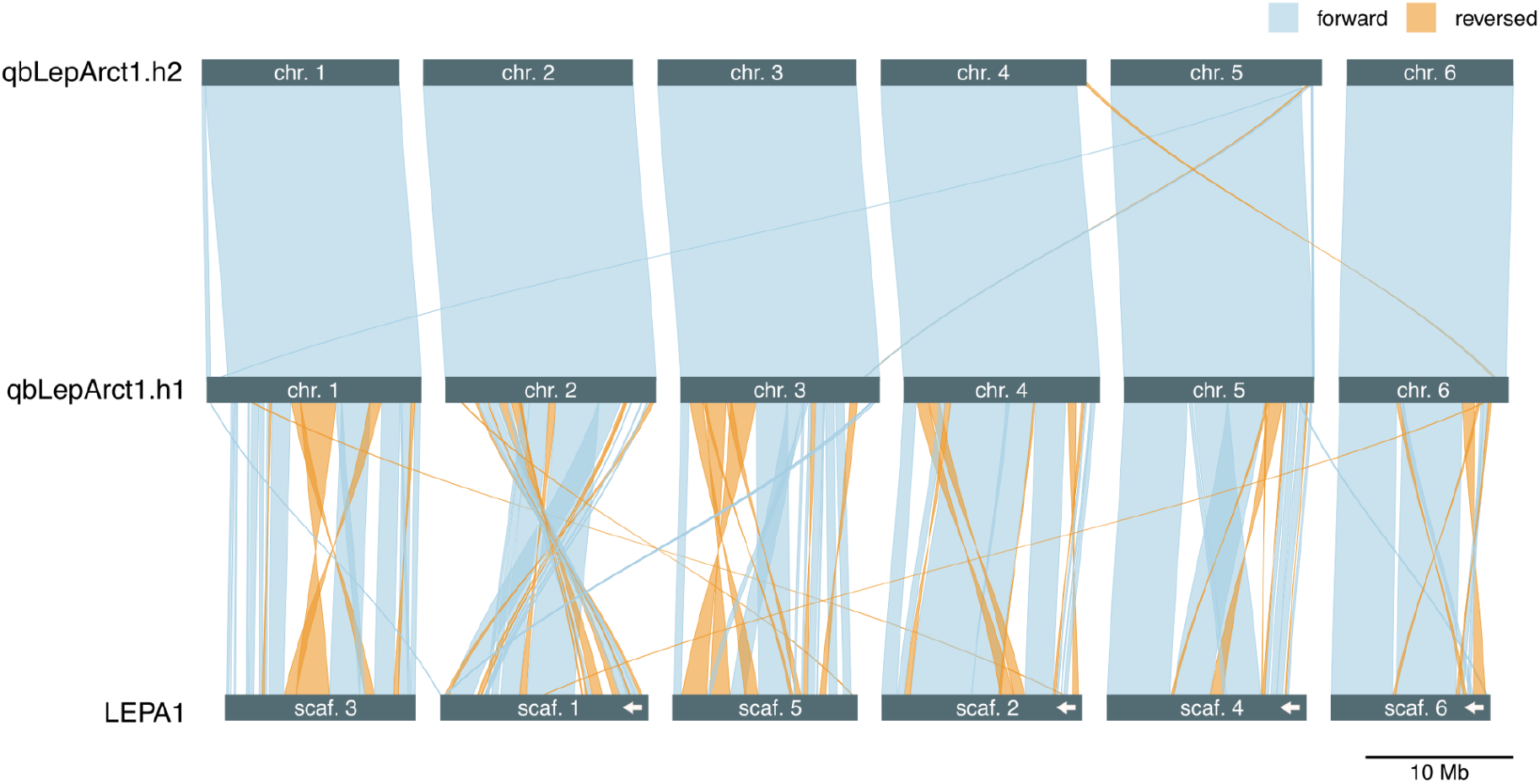
Whole-genome synteny between *Lepidurus packardi* (LEPA1) and the two *L. arcticus* haplotype assemblies, hap1 and hap2. Chromosomes are shown as horizontal tracks, arrows within LEPA1 scaffolds indicate reversals relative to the original FASTA orientation, and links indicate homologous aligned regions between adjacent assemblies. Only alignments with mapping quality at least 30 and alignment length at least 50 kb are shown. Chromosome order was anchored on hap1. Gray links denote the same orientation, whereas orange links denote reversed orientation. Scale bar, 10 Mb.

Total transposable element coverage was slightly higher in LEPA1 than in the two qbLepArct1 haplotypes (26.5% versus 24.1 - 24.9%; Supplementary Table 1). The clearest difference was in the fraction of nested and overlapping TE annotations. Nested TE coverage was 1.616% in LEPA1 compared with 0.180–0.181% in the qbLepArct1 haplotypes, while overlapping coverage was 0.905% compared with 0.054–0.065%. These differences were mainly driven by LTR and LINE elements, which contributed most of the nested and overlapping repeat coverage in LEPA1.

The repeat landscapes showed distinct Kimura distance profiles among assemblies (Fig. 5). Compared with the two *L. arcticus* haplotypes that are highly similar, *L. packardi* had a recent repeat peak at Kimura distance 0.0, corresponding to approximately 4% of the genome. It also showed a broad older repeat signal centred around 0.35–0.44, mostly confined to minor scaffolds. Around Kimura distance 0.1, the repeat signal in the qbLepArct1 haplotypes was mainly composed of LTR elements, whereas the corresponding signal in LEPA1 was dominated by LINE elements.

**Figure 5.**
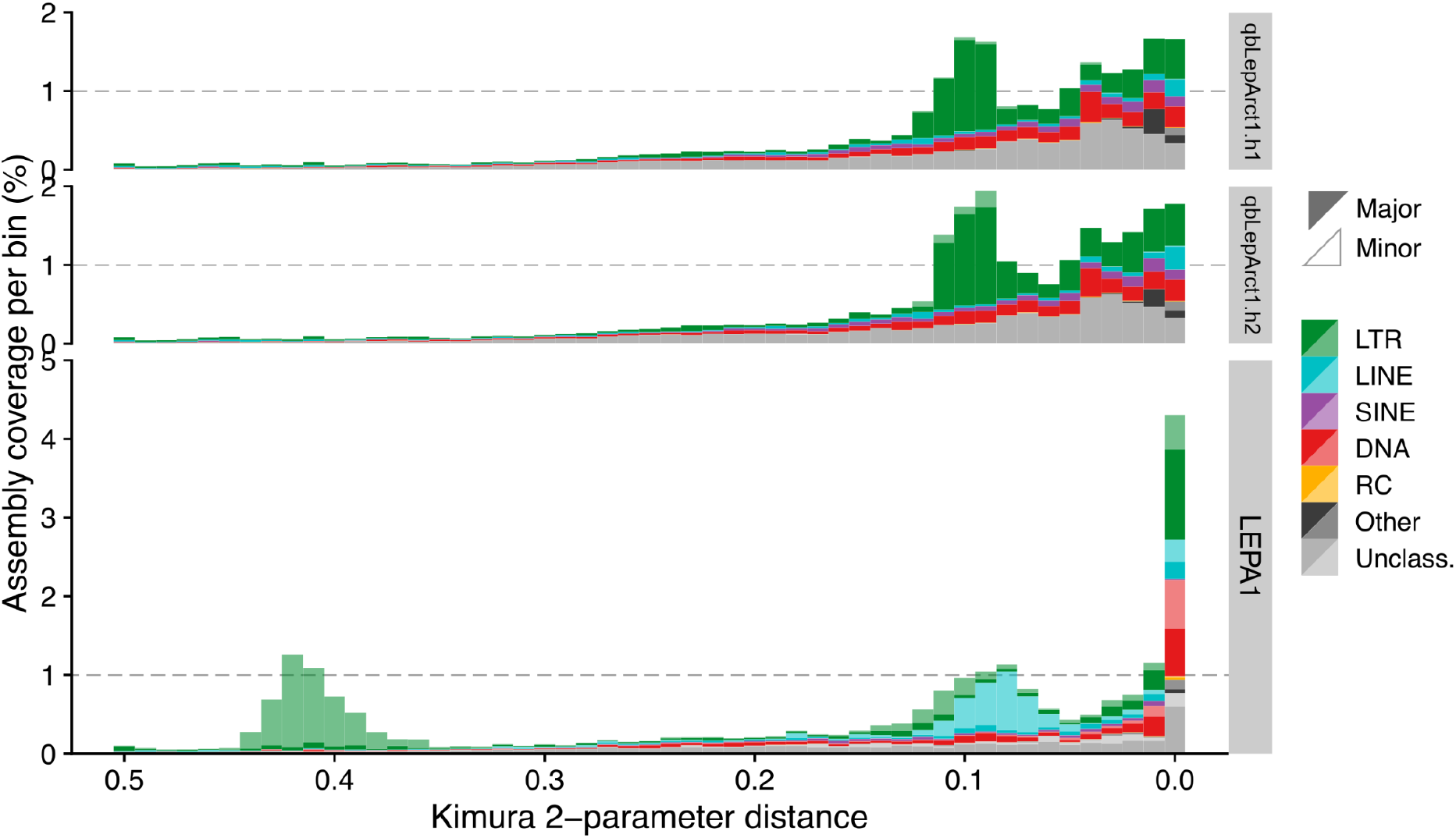
Transposable element age landscapes across assemblies. Kimura 2-parameter distance distributions of transposable element (TE) annotations in hap1 (qbLepArct1.1.hap1), hap2 (qbLepArct1.1.hap2), and LEPA1. Bars show assembly coverage per bin, and colors show TE classes: long terminal repeat retrotransposons (LTR), long interspersed nuclear elements (LINE), short interspersed nuclear elements (SINE), DNA transposons (DNA), rolling-circle transposons (RC), unclassified repeats, and other repeats. Opaque bars show repeats on the six largest chromosome-length major scaffolds, while transparent bars show repeats on smaller minor scaffolds. Lower Kimura distances indicate more recent TE copies.

Window-based TE summaries across the six largest scaffolds showed uneven TE distributions along chromosome-length scaffolds (Supplementary Fig. 3). TE-rich regions were often concentrated toward scaffold ends, particularly for LTR elements in the qbLepArct1 haplotypes. Similar LTR-rich terminal regions were present in some LEPA1 scaffolds, but LEPA1 also showed a larger contribution from LINE elements, with much of the LINE signal occurring on minor scaffolds not included among the six largest scaffolds.

## Discussion

Here we present the first chromosome-level reference genome for *Lepidurus arcticus*, a cold-adapted notostracan with a circumpolar distribution. The assembly process produced two pseudo-haplotype separated genome assemblies spanning 81.2 and 81.8 Mb, with more than 98% of each assembly assigned to six pseudo-chromosomes. Both assemblies were highly complete, with BUSCO completeness scores of 98.3%.

Hifiasm leverages the high accuracy of PacBio HiFi reads to preserve haplotype information in phased assembly graphs, enabling haplotype-aware assembly even at low heterozygosity (Cheng et al., 2021). Consistent with this, despite low estimated genome-wide heterozygosity of 0.133% (Fig. 1C; Supplementary Fig. 1), we recovered two highly complete pseudo-haplotypes, each with 98.3% BUSCO completeness and more than 95% HiFi k-mer completeness.

Comparison with *Lepidurus packardi* shows that the six main chromosomes are broadly conserved, however, with substantial intra-chromosomal rearrangements and inversions. The apparent intrachromosomal rearrangements are likely reflecting true structural divergence as expected from the two species evolutionary divergence. However, due to the less contiguous *L. packardi* assembly some of the divergence may also stem from assembly or scaffolding differences. Nevertheless, the high frequency of intra-chromosomal rearrangements between *L. arcticus* and *L. packardi* are not pointing in the direction of genetic evolutionary stasis, but rather point towards a normal evolutionary dynamics.

The TE landscapes suggest that repeat evolution in *L. arcticus* and *L. packardi* differs less by total TE content than by the timing and composition of repeat accumulation. Because Kimura divergence from the consensus can be used as a proxy for relative TE copy age (reviewed in (Rodriguez & Arkhipova, 2023), distinct peaks in these landscapes suggest differences in the timing of past repeat accumulation. *L. arcticus* shows a relatively consistent pattern across both haplotypes, with an intermediate burst largely associated with LTR elements. In *L. packardi*, the landscape is more heterogeneous, with a recent mixed-class peak, a LINE-associated intermediate burst, and an older LTR-associated burst not observed in *L. arcticus*. The greater nesting and overlap of LTR and LINE elements in *L. packardi* is consistent with this more complex age structure, suggesting repeated insertion into already repeat-rich regions. Together, these patterns suggest that *L. arcticus* has experienced a less complex history of repeat accumulation and turnover than *L. packardi*, although this interpretation should remain cautious because older repeat signals are harder to recover reliably and the *L. packardi* assembly is less contiguous.

Like other branchiopods, the small genome size (∼0.11 Gb) of *L. arcticus* is remarkable, and can possibly be linked to the low genetic variability and its phenotypic stasis over extended periods. Streamlined genomes among branchiopods is a shared phylogenetic trait (Alfsnes et al., 2017), and contrasts other arthropods like copepods that generally have large genomes and a flexible and complex life history (Leinaas et al., 2016; Schutt et al., 2021).

The two haplotypes of the reference genome display extremely low heterozygosity (0.133%). *Lepidurus arcticus* reproduces both sexually and asexually (Mathers, Hammond, Jenner, Zierold, et al., 2013; Weeks et al., 2006; Wojtasik & Bryłka™wołk, 2010). Under stable conditions (and low predation “stress”) clonal reproduction will dominate (Mathers, Hammond, Jenner, Zierold, et al., 2013). The sampled site is a pond devoid of fish and thus low predation risk, promoting clonal reproduction leading to the low heterozygosity.

*L. arcticus* is a “cold stenotherm” species, inhabiting cold alpine and arctic lakes and ponds. Cold habitats are often associated with large genomes (Hessen et al., 2013), but in the *Lepidurus* and *Triops* sister clades there seem to be no obvious patterns in this regard (Alfsnes et al., 2017). However, a full genomic or transcriptomic comparison between these warm- and cold-adapted clades *Triops* would be relevant to check for evolutionary adaptations to widely different temperature regimes.

This reference genome provides a basis for future comparative and population genomic studies of *L. arcticus*. It may help address questions about genome evolution, population connectivity, postglacial history, and adaptation in cold Arctic, subarctic, and alpine freshwater habitats.

## Acknowledgements

This project received data management and infrastructure support from ELIXIR Norway, supported by the Research Council of Norway’s grant 270068, the University of Bergen, the University of Oslo, the Arctic University of Norway in Tromsø, the Norwegian University of Science and Technology, and the Norwegian University of Life Sciences: NMBU. The authors acknowledge support from the National Infrastructure for High Performance Computing and resources provided by Sigma2 as well as Data Storage in Norway (project NN8013K) for computational work. The Norwegian Sequencing Centre generated the sequencing data used in this project (http://sequencing.uio.no).

## Data availability

Data generated for this study are available under ENA BioProject PRJEB111941. Raw PacBio sequencing data for the tadpole shrimp (ENA BioSample: SAMEA122213073) are deposited in ENA under ERX16434411, whereas Illumina Hi-C sequencing data are deposited in ENA under ERX16434412 and ERX16434413. Pseudo-haplotype one can be found in ENA at PRJEB111934, whereas hap2 is PRJEB111935.

Genome assemblies and gene annotations are also available at 10.5281/zenodo.19852443

## Supplementary

**Supplementary Figure 1.**
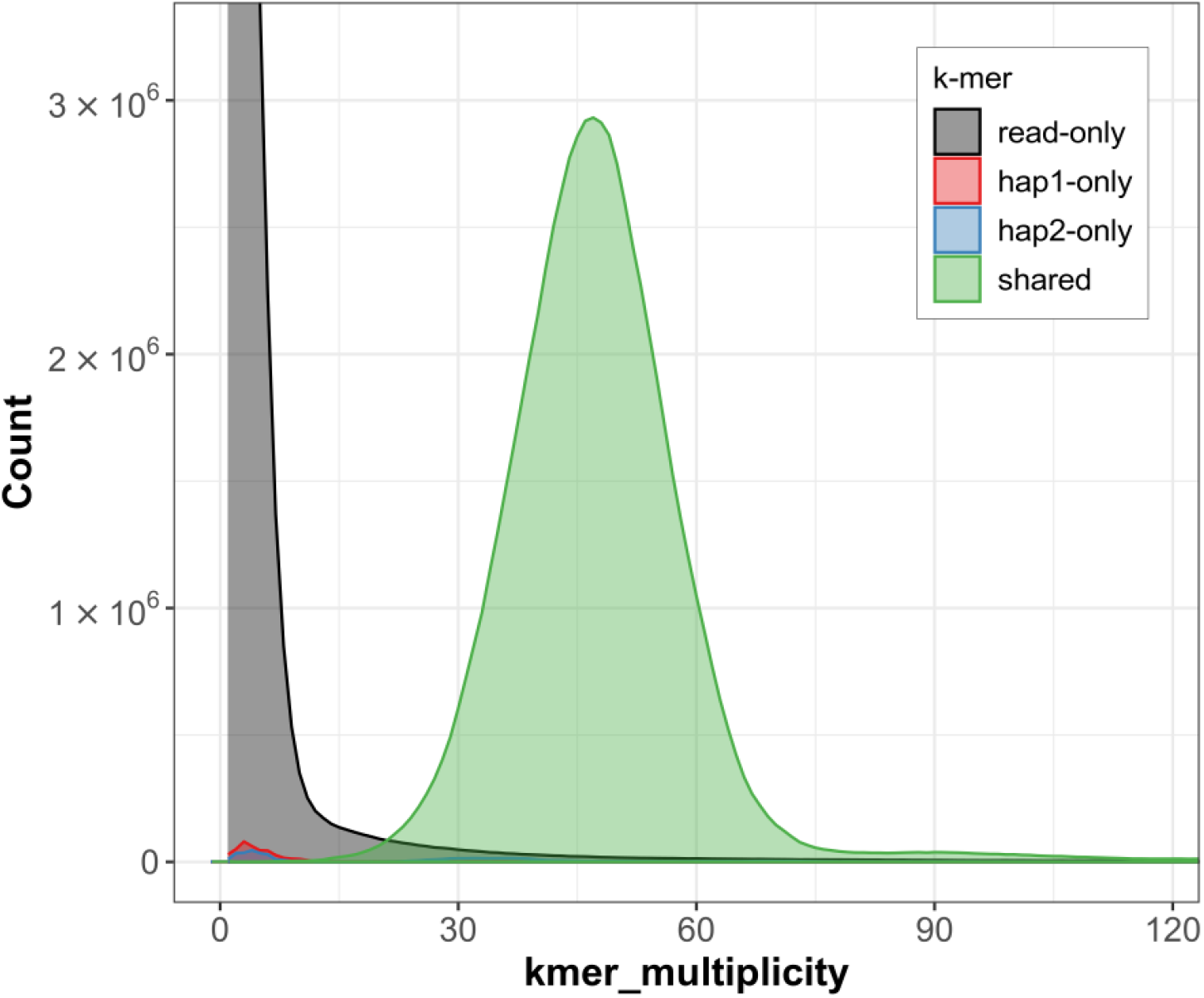
*K*-mer spectra of HiFi reads for *Lepidurus arcticus*. Distributions of *k*-mers found only in the reads (black), only in haplotype one (red), only in haplotype two (blue), or in both haplotypes (green). The y-axis represents the number of unique *k-*mers, while the x-axis represents the *k-*mer multiplicity (how often the *k*-mer is found in the set of HiFi reads).

**Supplementary Figure 2.**
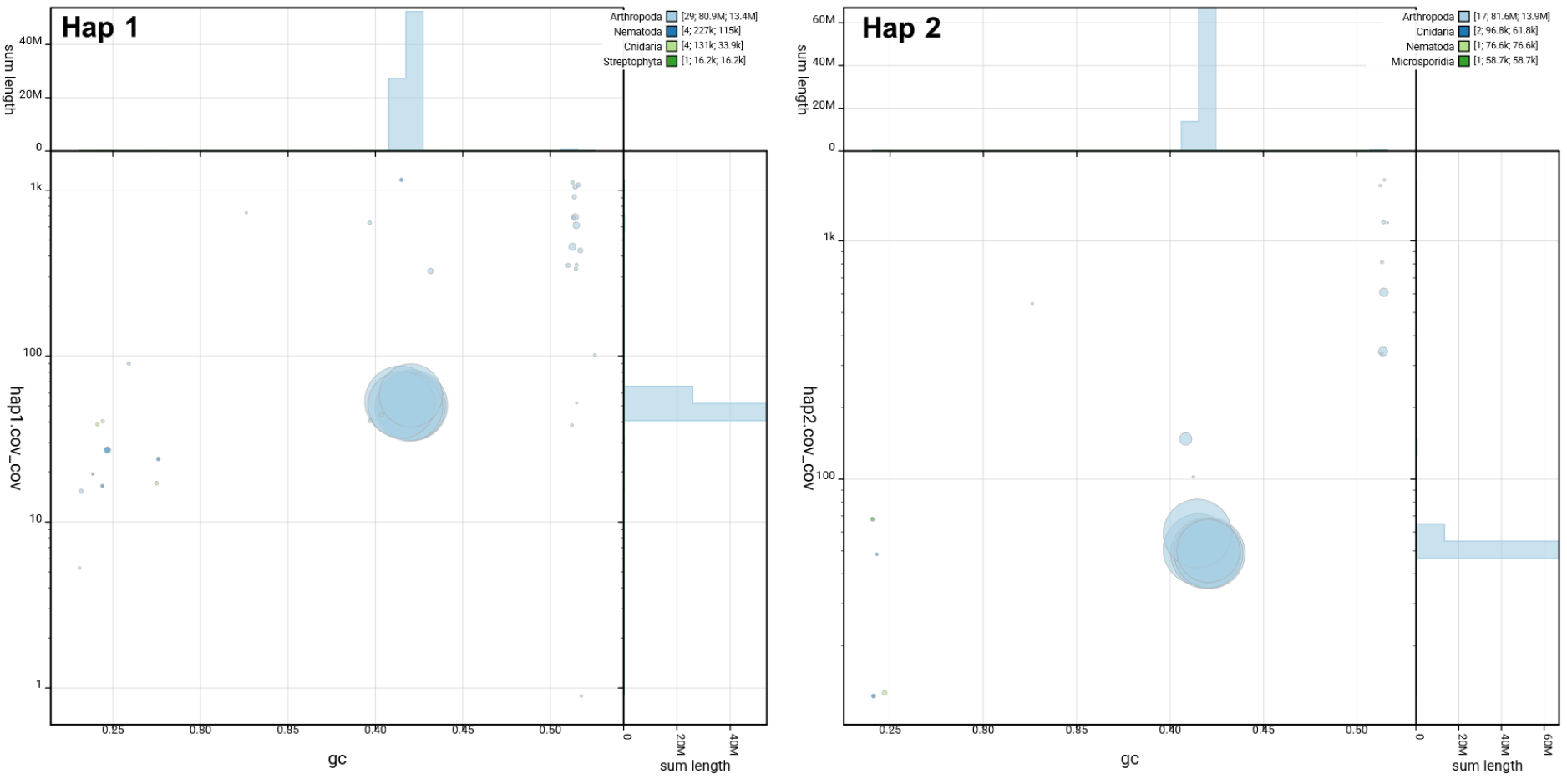
Coverage vs GC plots of *Lepidurus arcticus* hap1 and hap2. The BlobToolKit Blobplots depicts each scaffold as a dot based on the GC content (% GC, x-axis) and coverage (Y-axis). Size of the dots correspond to scaffold length. Dots are colored based on assigned taxonomy. Histograms of sequence lengths within a certain % GC range or coverage range are depicted on the top and right respectively.

**Supplementary Figure 3.**
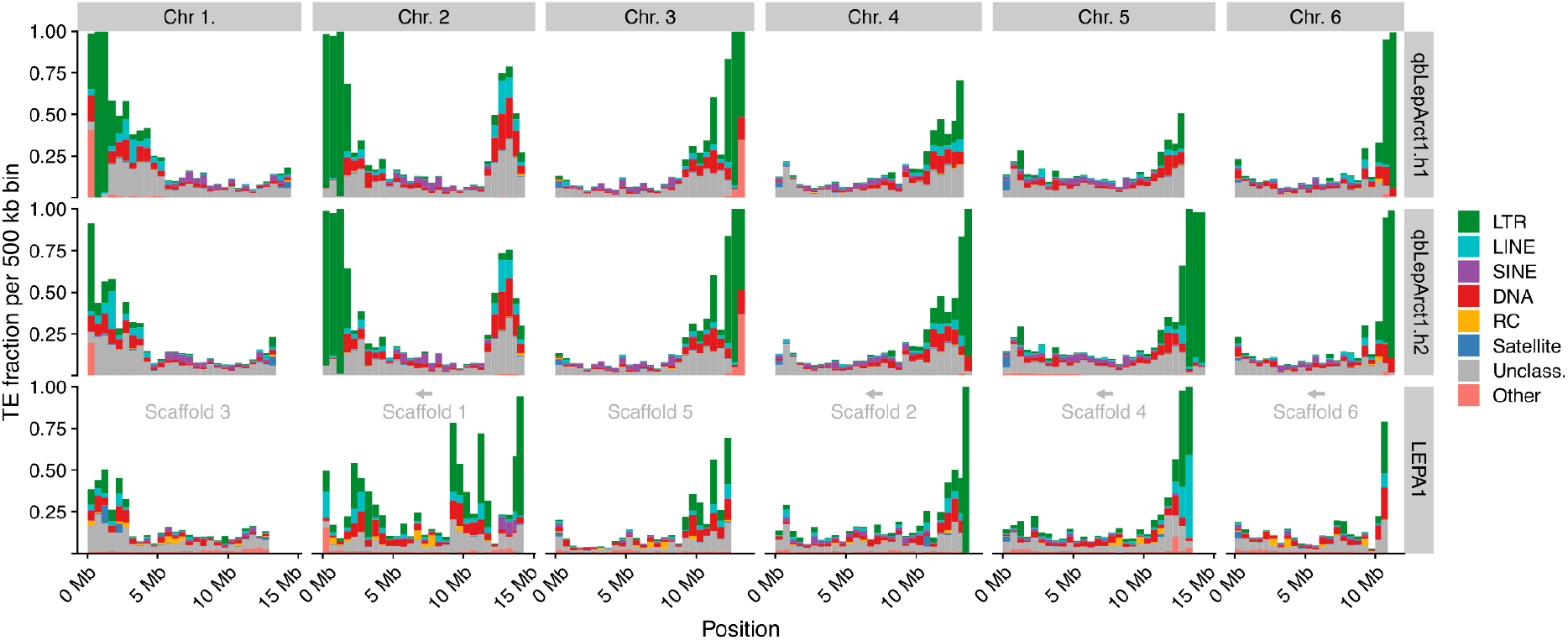
TE distribution across chromosome length scaffolds. Stacked TE density across the six largest chromosome-length scaffolds in hap1, hap2, and LEPA1, measured in 500 kb windows. Bar height shows the TE-covered fraction of each window, and colors show TE classes. Long terminal repeat retrotransposons (LTR), long interspersed nuclear elements (LINE), short interspersed nuclear elements (SINE), DNA transposons (DNA), rolling-circle transposons (RC), Satellite, Unclassified, and Other. Original scaffold names are shown in each LEPA1 panel with arrows indicating reverse complements relative to the original FASTA orientation.

**Supplementary Table 1.**
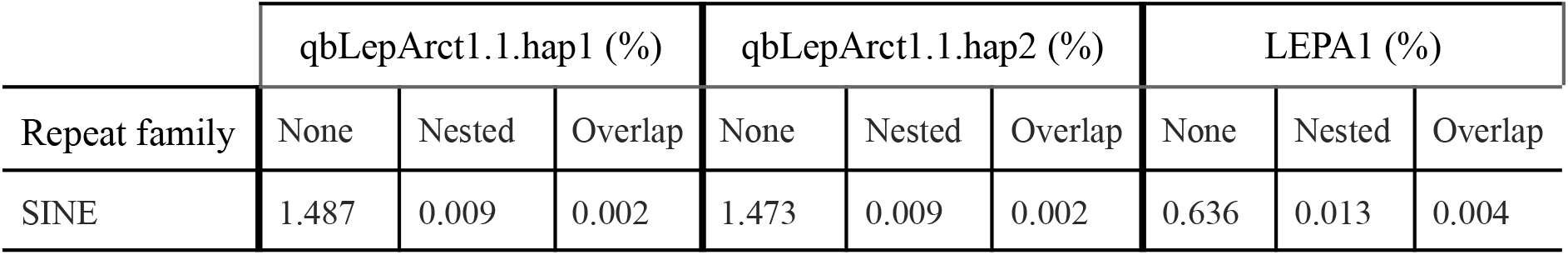
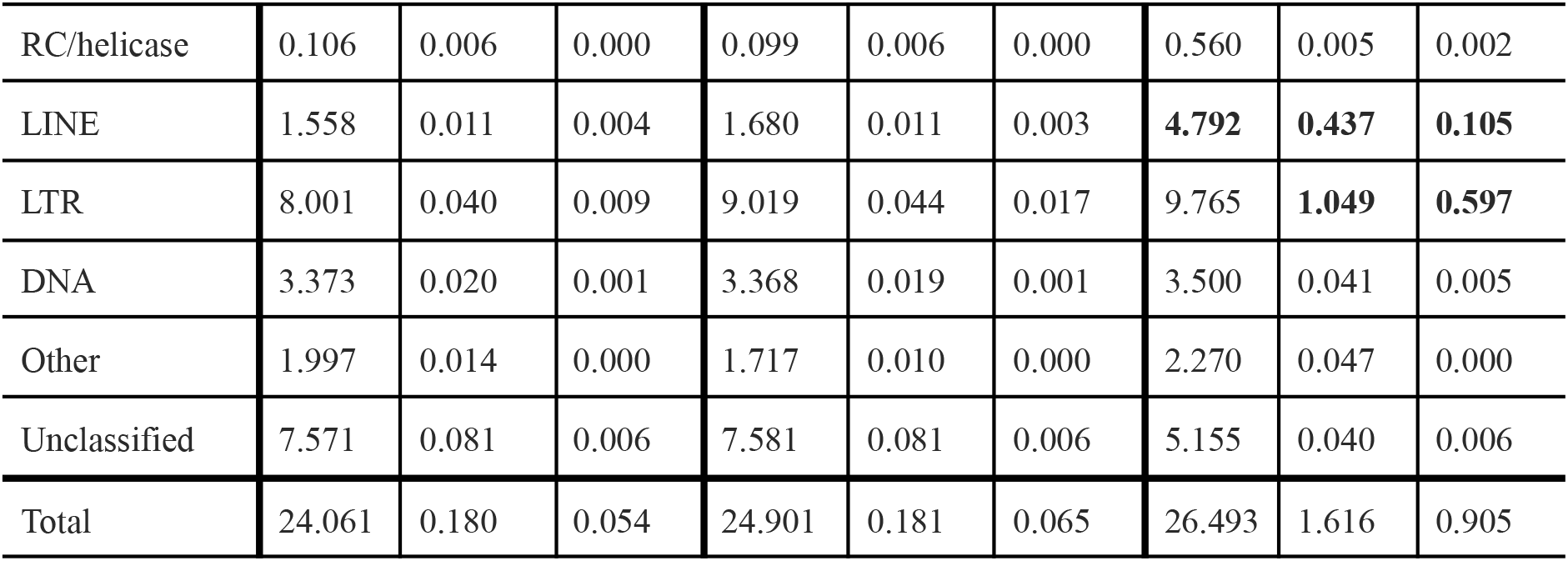
TE content in *Lepidurus arcticus* hap1, hap2, and the *Lepidurus packardi* LEPA1 assembly, shown as percentage of assembly length. The “None” TE coverage excludes fully nested annotations and overlapping annotation bases. Nested and overlap columns show the corresponding excluded fractions. Notable values of the differences are marked in bold.

