## Supplementary for "A chromosome-level genome assembly of “a living fossil”, the tadpole shrimp *Lepidurus arcticus* (Pallas, 1793)"

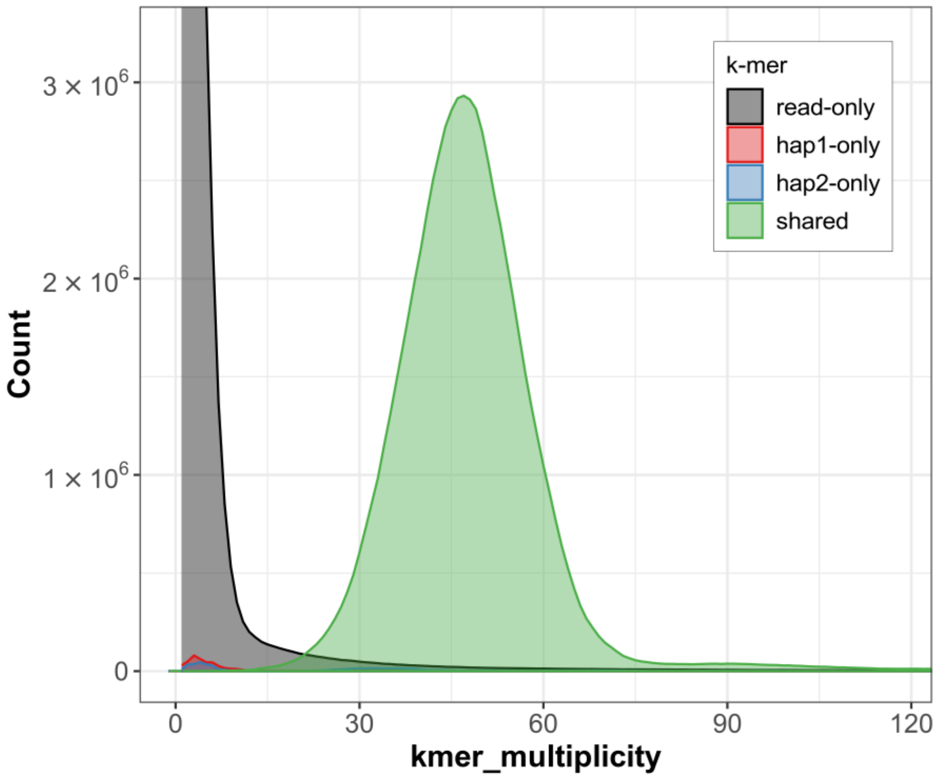


**Supplementary Figure 1. *K*-mer spectra of HiFi reads for *Lepidurus arcticus*.** Distributions of *k*-mers found only in the reads (black), only in haplotype one (red), only in haplotype two (blue), or in both haplotypes (green). The y-axis represents the number of unique *k*-mers, while the x-axis represents the *k*-mer multiplicity (how often the *k*-mer is found in the set of HiFi reads).

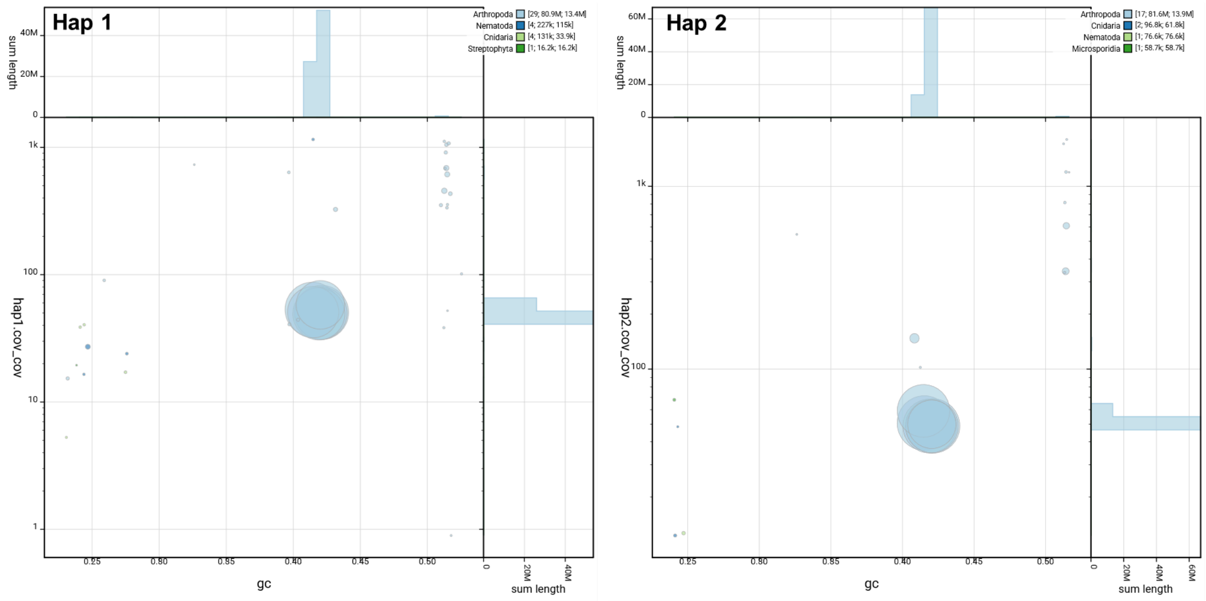


**Supplementary Figure 2. Coverage vs GC plots of *Lepidurus arcticus* hap1 and hap2.** The BlobToolKit Blobplots depicts each scaffold as a dot based on the GC content (% GC, x-axis) and coverage (Y-axis). Size of the dots correspond to scaffold length. Dots are colored based on assigned taxonomy. Histograms of sequence lengths within a certain % GC range or coverage range are depicted on the top and right respectively.

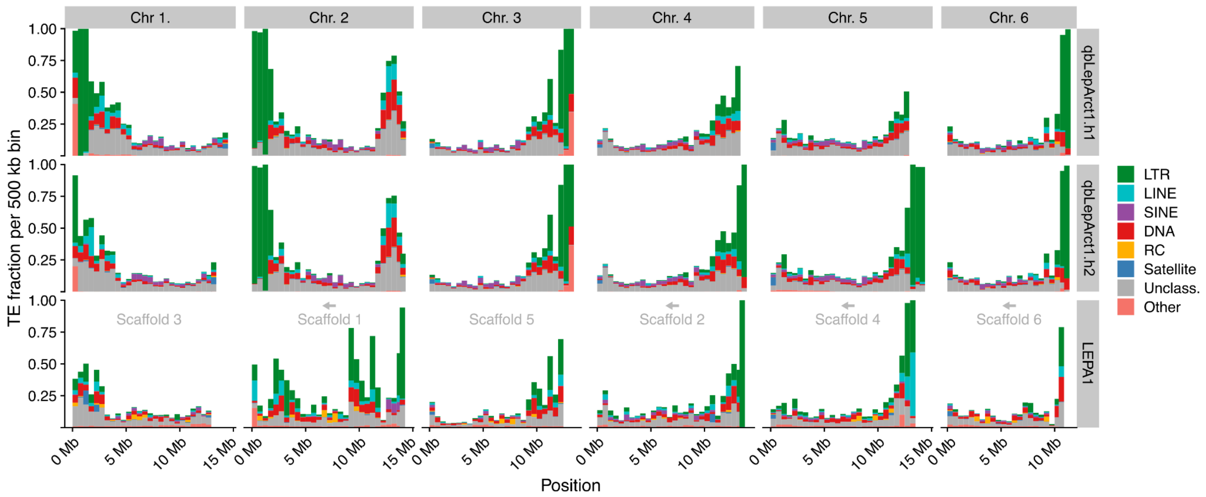


**Supplementary Figure 3. TE distribution across chromosome length scaffolds.** Stacked TE density across the six largest chromosome-length scaffolds in hap1, hap2, and LEPA1, measured in 500 kb windows. Bar height shows the TE-covered fraction of each window, and colors show TE classes. Long terminal repeat retrotransposons (LTR), long interspersed nuclear elements (LINE), short interspersed nuclear elements (SINE), DNA transposons (DNA), rolling-circle transposons (RC), Satellite, Unclassified, and Other. Original scaffold names are shown in each LEPA1 panel with arrows indicating reverse complements relative to the original FASTA orientation.

**Supplementary Table 1: TE content in *Lepidurus arcticus* hap1, hap2, and the *Lepidurus packardi* LEPA1 assembly, shown as percentage of assembly length.** The "None" TE coverage excludes fully nested annotations and overlapping annotation bases. Nested and overlap columns show the corresponding excluded fractions. Notable values of the differences are marked in bold.

| | qbLepArct1.1.hap1 (%) | | | qbLepArct1.1.hap2 (%) | | | LEPA1 (%) | | |
|---|---|---|---|---|---|---|---|---|---|
| Repeat family | None | Nested | Overlap | None | Nested | Overlap | None | Nested | Overlap |
| SINE | 1.487 | 0.009 | 0.002 | 1.473 | 0.009 | 0.002 | 0.636 | 0.013 | 0.004 |
| RC/helicase | 0.106 | 0.006 | 0.000 | 0.099 | 0.006 | 0.000 | 0.560 | 0.005 | 0.002 |
| LINE | 1.558 | 0.011 | 0.004 | 1.680 | 0.011 | 0.003 | **4.792** | **0.437** | **0.105** |
| LTR | 8.001 | 0.040 | 0.009 | 9.019 | 0.044 | 0.017 | 9.765 | **1.049** | **0.597** |
| DNA | 3.373 | 0.020 | 0.001 | 3.368 | 0.019 | 0.001 | 3.500 | 0.041 | 0.005 |
| Other | 1.997 | 0.014 | 0.000 | 1.717 | 0.010 | 0.000 | 2.270 | 0.047 | 0.000 |
| Unclassified | 7.571 | 0.081 | 0.006 | 7.581 | 0.081 | 0.006 | 5.155 | 0.040 | 0.006 |
| Total | 24.061 | 0.180 | 0.054 | 24.901 | 0.181 | 0.065 | 26.493 | 1.616 | 0.905 |
